# Effects of Long-Term Meditation Practices on Sensorimotor Rhythm Based BCI Learning

**DOI:** 10.1101/2020.09.09.290080

**Authors:** Xiyuan Jiang, Emily Lopez, James Stieger, Carol Greco, Bin He

**Author notes:** **Correspondence:** Bin He.

## Abstract

Sensorimotor rhythm (SMR) based brain-computer interfaces (BCIs) provide an alternative pathway for users to perform motor control using motor imagery (MI). Despite the non-invasiveness, ease of use and low cost, this kind of BCI has limitation due to long training times and BCI inefficiency— where a subpopulation cannot generate decodable EEG signals to perform the control task. Meditation is a mental training method to improve mindfulness and awareness, and is reported to have a positive effect on one’s mental state. Here we investigate the behavioral and electrophysiological differences between experienced meditators and meditation naïve subjects in 1-dimensional and 2-dimensional cursor control tasks. We found that within subjects who have room for improvement, meditators outperformed control subjects in both tasks, and there were fewer BCI insufficient subjects in the meditator group. Finally, we also explored the neurophysiological difference between the two groups, and showed that meditators had higher SMR predictor and were better able to generate decodable EEG signals to achieve SMR BCI control.

## 1 Introduction

Decades of research have sought to find alternative methods of communication between the human brain and the outside world. With the ever-growing knowledge in the neuroscience field, scientists have designed the brain-computer interface (BCI) to achieve this goal (Wolpaw et al., 2002; He et al., 2020). A BCI attempts to recognize the user’s intent by decoding her/his neurophysiological signals and then converts this intent into commands to control objects, such as a cursor on a computer screen (Wolpaw et al., 1991; Trejo et al., 2006), a quadcopter (LaFleur et al., 2013) or a robotic arm in space (Meng et al., 2016; Edelman et al., 2019).

One of the main goals for the BCI is to help people suffering from various kinds of neuromuscular diseases, such as amyotrophic lateral sclerosis (ALS), stroke, and spinal cord injury (Armour et al., 2016), to regain a certain degree of movement ability (Rebsamen et al., 2010; Ang et al., 2015). Despite limited ability to move, cognitive ability in this population remains partially or fully intact. Therefore, it would be a significant improvement in quality of life to use a BCI to complete daily life tasks.

BCIs have been designed to decode various kinds of brain signals: such as neurons’ action potentials using multielectrode arrays (Maynard et al., 1997), electrical signal over the cortex using ECoG (Schalk et al., 2007), and electrical signal over the scalp using EEG (He et al., 2015). Among all the recording techniques, BCI based on EEG is one of the most widely used in research and clinical settings due to its ease of use, relatively low costs, and high temporal resolution (He et al., 2015). The SMR or mu rhythm is generated by the synchronized electric brain activity over the motor cortex area, and has a frequency range of around 8 - 12 Hz (Pfurtscheller et al., 2006; Bernier et al., 2007). In BCI application, the frequency band centered at 12Hz (Cassady et al., 2014; Meng et al., 2016, 2018; Stieger et al., 2020) was shown to be effective in SMR control. Event-related desynchronization (ERD) occurs when the amplitude of mu rhythm decreases in response to a person moving or imagining moving her/his body (Pfurtscheller and Aranibar, 1979). On the other hand, when a person stops moving or imaging moving, the amplitude of mu rhythm increases, termed event-related synchronization (ERS). The SMR based BCI is a well-established BCI modality, and it has been demonstrated that people can perform multi-dimensional cursor control (McFarland et al., 2010; Meng et al., 2018), drone control (LaFleur et al., 2013), wheelchair control (Galán et al., 2008; Huang et al., 2012), and robotic arm control (Meng et al., 2016; Edelman et al., 2019) with SMR BCI.

Despite the progress of SMR based BCI, challenges exist. For example, unlike EEG BCI based on P300 (Fazel-Rezai et al., 2012) and steady state visually evoked potentials (ssVEP) (Bakardjian et al., 2010), SMR based BCI usually requires several sessions of training, and around 20% of subjects are not able to achieve accurate control even after training (Blankertz et al., 2010). While efforts have been mainly focused on developing better decoding algorithms and recording techniques (Lotte and Guan, 2011), i.e. from the ‘computer’ perspective of BCI, limited attention has been drawn to enhancing people’s ability to generate more decodable EEG signals, i.e. from the ‘brain’ side. For the latter, the high-level goal is to determine, given the same BCI system, if there exists a subpopulation who is better able to control it, and if a certain kind of training or intervention could be developed to equip ordinary people with this BCI control ability.

Prior literature has suggested that meditation shows a distinct change in one’s brain structure, and meditators tend to develop the ability to better control their attention and awareness (Chan and Woollacott, 2007; Tang et al., 2007; Moore and Malinowski, 2009). In the search for optimal training methods in preparation for the SMR based BCI control, previous work has investigated whether people with meditation experience are better able to control SMR based BCI (Cassady et al., 2014; Tan et al., 2014, 2015; Kober et al., 2017; Stieger et al., 2020), or just generate ERD/ERS without controlling a BCI system (Kerr et al., 2013; Rimbert et al., 2019). Similar to what Tang and colleagues (Tang et al., 2015) summarized for the neuroscience aspect of meditation studies, efforts to study the meditation effect on SMR BCI could be divided into two categories, longitudinal studies and cross-sectional studies:

1. Longitudinal studies separated meditation-naïve subjects into a meditation group and a control group, with the meditation group receiving meditation training and control group receive either active control tasks or no specific task (Tan et al., 2014, 2015; Botrel and Kübler, 2019; Stieger et al., 2020). After that, BCI performance and/or neurophysiological difference between the two groups was assessed.
2. Cross-sectional studies investigated the difference in BCI/neurofeedback learning between people who already have meditation experience and meditation naïve subjects (Cassady et al., 2014; Kober et al., 2017). In Cassady and colleague’s work (Cassady et al., 2014), the meditation group was shown to have better performance compared with the control group in terms of performance, learning speed, and information transfer rate. However, most of the claims in this study focused on the behavior difference. A more in-depth analysis of the neurophysiological difference is needed. Another question left unanswered is whether meditators are also better at more complex tasks, such as 2-dimension (2D) cursor control. In another study, Kober and colleague (Kober et al., 2017) found that people who pray frequently had a higher ability to control the SMR, but the recording was limited to Cz electrode only and the control dimension was limited to 1-dimension (1D).

Is meditation experience indeed a significant factor affecting SMR BCI learning? For example, Stieger and colleagues (Stieger et al., 2020) found that after an 8-week mindfulness-based stress reduction training, subjects indeed had significant performance improvements in the up/down task (both hands motor imagery to go up and rest to go down), but for the left/right control task (left/right hand motor imagery) the effect was not significant. Botrel and colleagues (Botrel and Kübler, 2019) found that week-long visuomotor coordination and relaxation training does not improve the SMR based BCI performance. One of the reasons for this kind of disagreement may be a dose-effect, meaning that it might take a longer meditation time to affect BCI learning in a significant manner.

With these questions in mind, we recruited experienced meditators and controls and investigated the difference in SMR BCI learning between these two groups in both 1D and 2D tasks. The aims for this cross-sectional study are as follows: First, to verify the conclusions in the pilot study (Cassady et al., 2014) that meditators had better learning in SMR BCI with an independent investigation; Second, to explore the behavior difference between the two groups in a more complex 2D task; Third, to investigate the neurophysiological difference between these two groups.

## 2 Methods

### 2.1 Participants

The experimental procedures involving human subjects described in the current study were approved by the Institutional Review Board (IRB) of Carnegie Mellon University, all participants provided written informed consent. We utilized a single-blind two-group experimental design, with a meditation group and a control group. The experimenters did not know the identity of the subject in relation to their meditation experience throughout the whole study, and avoided any conversation related to meditation. Subjects were recruited via flyers in the surrounding area, as well as an email sent out to local mindfulness groups. The meditation group consisted of 16 healthy subjects (age = 37.6 +/− 15.1) with a history of meditation practice, as evaluated by a questionnaire regarding personal meditation practice completed prior to experimentation. To be accepted into the meditator group, individuals had to cite at least a year of frequent and consistent practice, with most subjects having 2 or more years of consistent practice. Most of the meditators’ practices belong to the subgroup of Vipassana, Zen, Mindfulness, and Buddhism. The control group consists of 19 healthy individuals (age = 24.8 +/− 8.7) with no prior meditation experience. Both groups had no prior BCI experience. We continually asked participants to describe their motor imagery strategies. If these strategies diverged from the kinesthetic motor imagery they were asked to perform, we reminded them to focus on the sensations and intention behind the imagined motion of their hands. We excluded one subject (identity: meditator) from the analysis because she/he did not follow the motor imagery guidelines.

### 2.2 Surveys

In the first session, we asked subjects to fill out two surveys before the BCI experiment. Both surveys aim to measure one’s level of mindfulness. The first survey is called the Freiburg Mindfulness Inventory (FMI) (Walach et al., 2006), which has 14 statements such as ‘I am open to the experience of the present moment’. The subject was asked to use a 1-4 scale to indicate how often she/he has such experience. The FMI score was calculated by summing up the answers to each question with proper recode of one question (Walach et al., 2006). The second survey is called Day-to-Day Experiences (Brown and Ryan, 2003), which has 15 questions, such as ‘I find it difficult to stay focused on what’s happening in the present’, the subject was asked to use a 1-6 scale to indicate how often she/he has such experience. In the end, the Mindful Attention Awareness Scale (MAAS) was calculated by averaging answers to each question. In both surveys, higher score indicates higher level of mindfulness.

### 2.3 Data acquisition

Subjects in both groups went through 6 sessions of BCI training within 4 – 6 weeks, with at least one experiment per week. Each experimental session lasted about 2 hours, with a 9-minute break in the middle. EEG data were recorded throughout the session using the Neuroscan SynAmps system with 64-channel EEG QuikCap (Neuroscan Inc, Charlotte, NC). The sampling frequency was set to 1000 Hz, and the impedance was kept below 5kΩ during the preparation. The experimenter checked the impedance in the break to make sure it remained below 5kΩ.

The experiment setup is shown in Figure 1. Each session began with a 5-minute warmup task, where the subject was instructed to perform left-or right-hand motor imagery by focusing on imagining the sensations and intention of opening/closing the left/right hand. After that, the subject was asked to perform BCI cursor control of three different tasks: left/right (LR), up/down (UD), and 2D, by moving the ball to the corresponding bar with motor imagery. BCI2000 was used to perform a standard SMR BCI cursor task (Schalk et al., 2004), where the mu rhythm band power of C3 and C4 electrodes after small Laplacian filter was used as features. In this work, the mu rhythm was set to be centered at 12 Hz (Meng et al., 2016, 2018) with a 3 Hz bin (Stieger et al., 2020), and was estimated using an autoregressive approach. In the LR task, subjects were told to image opening/closing the right hand as they practiced in the warm-up to move the ball to the right, and left-hand motor imagery to move the ball to the left. After subjects performed 3 rounds of LR BCI, with each round consisting of 25 trials, a similar explanation was given for the up-down (UD) BCI task, except they were instructed to imagine both hands opening and closing to move the ball up, and to rest, in other words try to clear their minds to move the ball down. After subjects performed three rounds of UD BCI, they moved onto the 2D task, in which the same instructions were implemented to move the ball up, down, left, or right according to which bar appeared on the screen. After one block (3 rounds) each of LR, UD, and 2D BCI, the subjects were given a 9-minute break in which they were instructed to read and rate comics by pressing a key on the keyboard, this standard ‘break task’ ensures that subjects use the same approach to relax. After the break, they completed one more block each of LR, UD, and 2D BCI.

**Figure 1.**
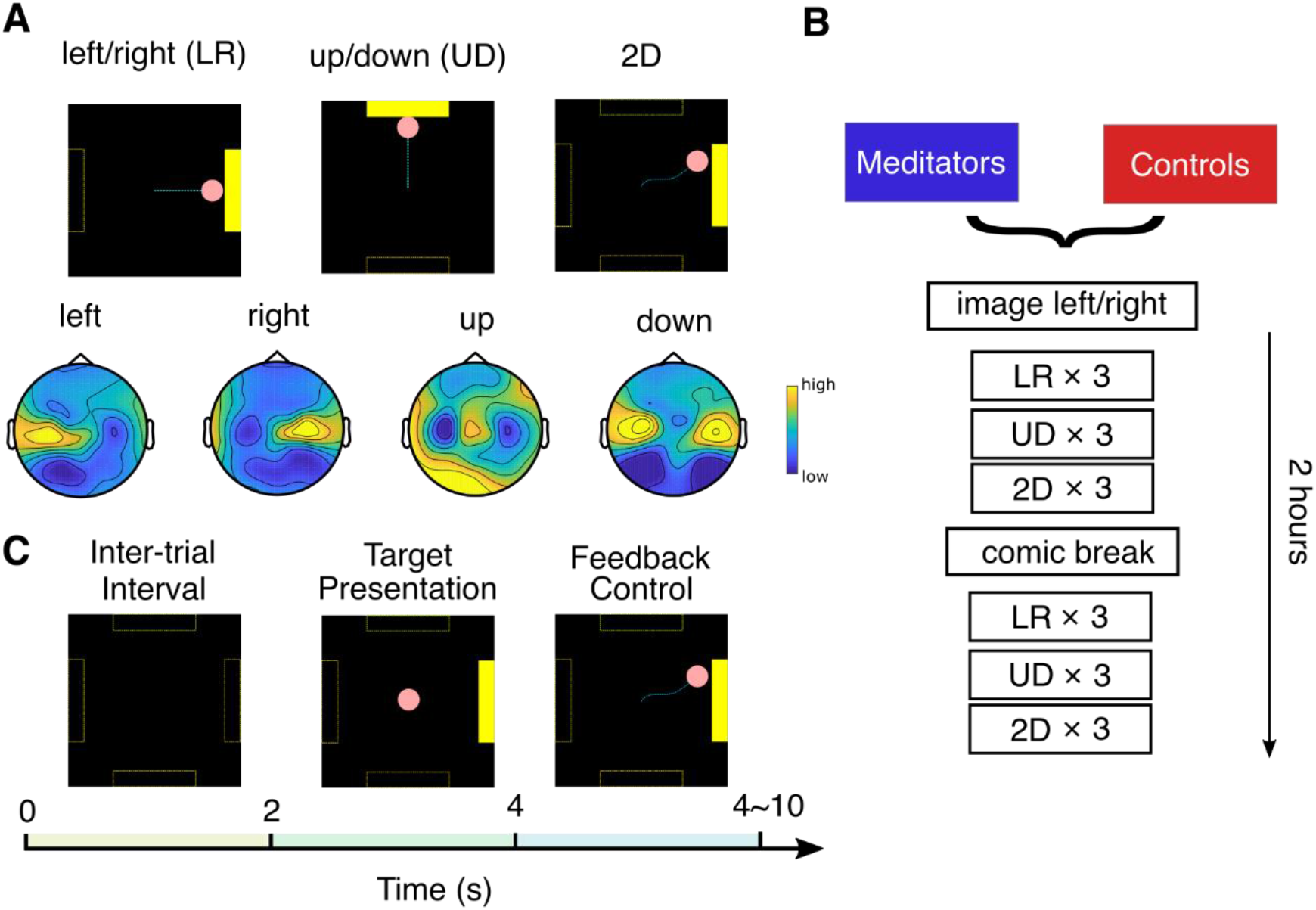
Experimental setup. (**A)** Top: three experiment tasks and typical cursor trajectories in left/right (LR) control, up/down (UD) control and 2D control. The dashed lines were invisible to the subject. Bottom: example topology of mu rhythm band power in each motor imagery class. **(B)** Experiment flow of one session. **(C)** Each trial consists of 2s of inter-trial interval, 2s of target presentation and 0~6s of BCI feedback control.

### 2.4 Performance metric

We quantify the performance using percent valid correct, or PVC (Cassady et al., 2014; Meng et al., 2016; Edelman et al., 2019), which is the ratio between the number of hit trials and number of hit trials plus the number of missed trials. When analyzing the data, unless stated otherwise, we excluded subjects with baseline PVC > 90% in both LR and UD conditions, because these subjects usually did not have much room for learning. 4 control subjects were excluded under this criterion, accounting for 11% of the total subjects. Together with the subject excluded due to not following the MI guideline (1 meditator), the number of subjects involved in the analysis is 15 meditators and 15 controls.

### 2.5 Offline EEG data analysis

We bandpass filtered the EEG data using a Hamming window sinc FIR filter with the passband set between 1 Hz and 100 Hz, then down sampled to 250Hz. ICA was performed to remove artifacts such as eye blinking. After that, complex Morlet wavelet convolution was used to extract the power of the mu frequency band (3 Hz bin centered at 12 Hz).

The neurophysiological predictor, or SMR predictor measures the difference between mu band power and the 1/*f* noise floor in a power-frequency plot for C3 and C4 (Blankertz et al., 2010). Concretely, the EEG power spectrum at rest could be fitted with the sum of a 1/*f* noise floor, *n*(*f*; *λ*, ***k*** _*n*_) and two Gaussian distributions, centered at mu rhythm and beta rhythm, *g*_*α*_(*f*; *μ*_*α*_, *σ*_*α*_) and *g*_*β*_(*f*; *μ*_*β*_, *σ*_*β*_). In this study, the power spectral density is equal to the mean of C3 and C4 band power after small Laplacian spatial filtering during the inter-trial resting state, combining LR conditions and UD conditions.

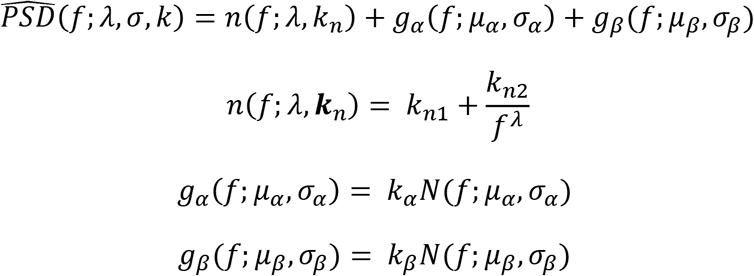

The SMR predictor (dB) is calculated individually for C3 and C4 electrode mu rhythm band power after small Laplacian spatial filtering.

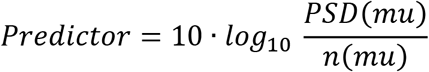

In the case where the algorithm could not find a curve to fit, we manually selected 5~10 representative data points to describe the 1/*f* noise floor function by following the trend of the PSD curve and fitted these points using *n*(*f*; *λ*, ***k***_*n*_). We discard a subject and session pair if the PSD does not follow a 1/*f* decrease trend. The percentage of data points discarded was 10.5%.

We designed a method to calculate the control signal during task execution to be as close to the real condition as possible. Concretely, we first calculated the C3 and C4 electrode frequency band power after small Laplacian spatial filtering, denoted *P*_*C*3_ and *P*_*C*4_. Then the raw control signal was calculated using the following equation:

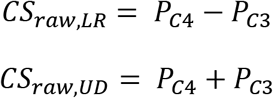

Then we applied a similar z-scored procedure to the raw control signal as the BCI 2000 platform,

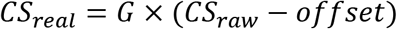

Where *G* and *offset* are set to make the *CS*_*real*_ zero mean and unit variance. The difference between this offline z-score and online approach is that the latter is causal and adaptive, i.e. *G* and *offset* is calculated via past 30 seconds of window, and change as time goes on. As shown in Supp Figure S1, we found that control signal under this definition could better explain the variability of performance than the ERD/ERS method, i.e. band power during task execution divided by resting state band power.

We quantify the contrast between two contexts in a task (e.g. left trials and right trials in LR task) using Fisher score (Perdikis et al., 2018).

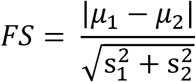

Where *μ*_1_ and *μ*_2_ are the means and 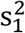 and 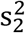 are the variance of context 1 and context 2’s band power in one session. The fisher score is calculated independently for each channel.

## 3 Results

### 3.1 Survey results

In both surveys, we found meditators had higher scores than control subjects. Concretely, the FMI score for meditator is 45.2 ± 5.0, while for control subject it is 37.3 ± 6.9. The difference is significant (Wilcoxon rank-sum test, Z = 3.11, p = 0.0018). The MAAS score for meditator is 4.51 ± 0.84, while for control subject it is 3.74 ± 0.67. The difference is significant (Wilcoxon rank-sum test, Z = 2.69, p = 0.007). Bar plots for the two groups’ scores are shown in Figure 2(A). The same observation also holds when including subjects who are BCI proficient at baseline. These results serve as an additional proof, apart from the self-reported meditation experiences, that the meditators had higher level of mindfulness than the control group. In addition to the group difference, we also calculated the correlation between these survey results and performance. We used baseline PVC as performance because this session is when the surveys were filled out. The correlation between survey results and UD PVC turned out to be significant. Specifically, for FMI, r = 0.41, p = 0.014, and for MAAS, r = 0.41, p = 0.017.

**Figure 2.**
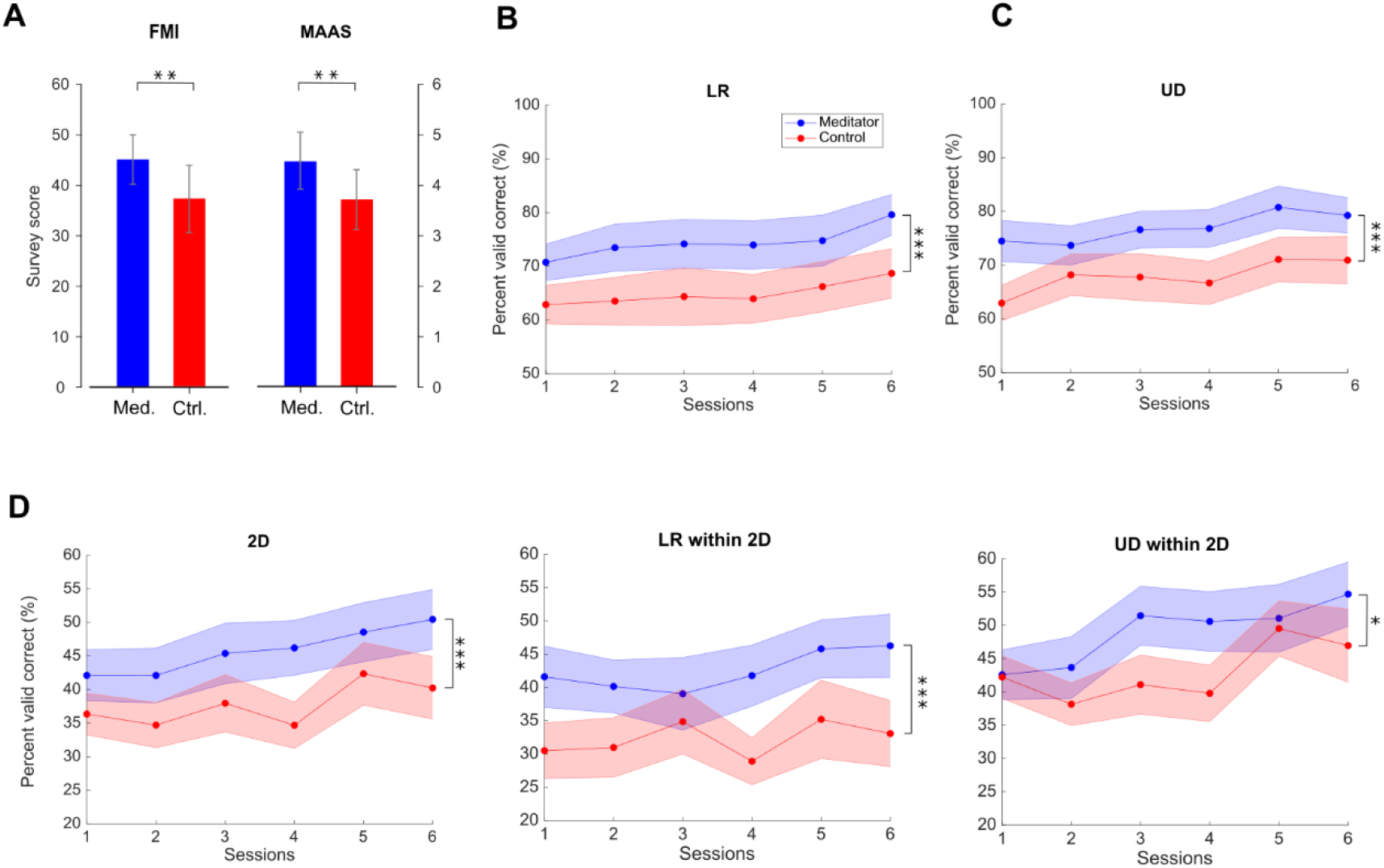
Survey results and group averaged performance and learning. **(A)** Survey results of FMI and MAAS shows that meditators have higher level of mindfulness than controls. Data are shown as mean±SD. Med. is meditator, Ctrl. is control. **(B)** Group LR averaged PVC±SEM for mediators and controls. Asterisk indicates significant group effect between meditator and controls with ANCOVA with p < 0.05, **(C)** for UD task, (d) for 2D task, LR within the 2D task and 2D within the 2D task. * indicates group difference with p < 0.05, ** indicates p < 0.01, *** indicates p < 0.001, same for subsequent plots.

### 3.2 Group averaged performance

Within the population who are not BCI proficient at baseline (i.e. subjects did not have > 90% of PVC in both LR and UD in session 1), we found that meditators achieved better performance (PVC) compared with control subjects, and this difference was consistent throughout the six sessions. The group averaged performance in the baseline and final session is shown in Table S1, and the averaged performance for all sessions is shown in Figure 2.

We modeled the learning progress as a linear regression model. To see if the regression lines between meditators and controls are different, we used analysis of covariance (ANCOVA). We found that the difference between groups was significant in all three tasks (F(1,176) = 14.62, 17.34, 12.16 with p = 0.0002, <0.0001, 0.0006 for LR, UD and 2D). The learning effects in UD and 2D task were also significant (F(1,176) = 4.38, 4.51, p = 0.03, 0.03 for UD and 2D). However, the learning effect did not show significance in LR (F(1,176) = 2.76, p = 0.10). There were no significant interaction (session x group) effects in any of the three tasks (F(1,176) = 0.05, 0, 0.23 with p = 0.83, 0.96, 0.06 for LR, UD and 2D), indicating that the learning speed was not different between two groups.

Realizing the fact that the 2D task is the combination of LR and UD, we next separated the LR and UD task within the 2D. Interestingly, we found that within the 2D task, meditators had a higher baseline of LR, but for the UD these two groups were at the same level. Further, the learning curve showed that meditators had numerically better learning compared with controls in the UD within 2D. Statistical analysis using ANCOVA shows that performance in both LR and UD within the 2D task was different between two groups (F(1,176) = 14.83, 5.91, p = 0.0002, 0.016 for LR and UD within 2D), as well as the learning effect of UD (F(1,176) = 7.36, p = 0.0007). On the other hand, LR within 2D did not show a significant learning effect (F(1,176) = 1.33, p = 0.25). The learning speed between two groups was not significantly different between groups as well (F(1,176) = 0.19, 0.25, p = 0.665, 0.619 for LR and UD within 2D).

### 3.3 Competency curve

While group averaged PVC is a good indicator of performance, there are several drawbacks. First, it only provides information on the overall trend of performance during BCI learning; we still do not know how many subjects remain BCI inefficient. Second, it does not provide information regarding within-session learning.

To intuitively show how learning occurs in the two groups, we plotted the percentage of subjects whose PVC remained below a threshold, as sessions go on. We set the threshold as 70% for 1D control and 40% for 2D control (Combrisson and Jerbi, 2015), but we obtain similar results under varied thresholds. To cope with potential fluctuation of performance, a subject passes the threshold if he/she meets one of the following criteria: achieving an averaged PVC > threshold in three consecutive runs, or achieving an averaged PVC > threshold in one single session (Cassady et al., 2014). The result is shown in Figure 3.

**Figure 3.**
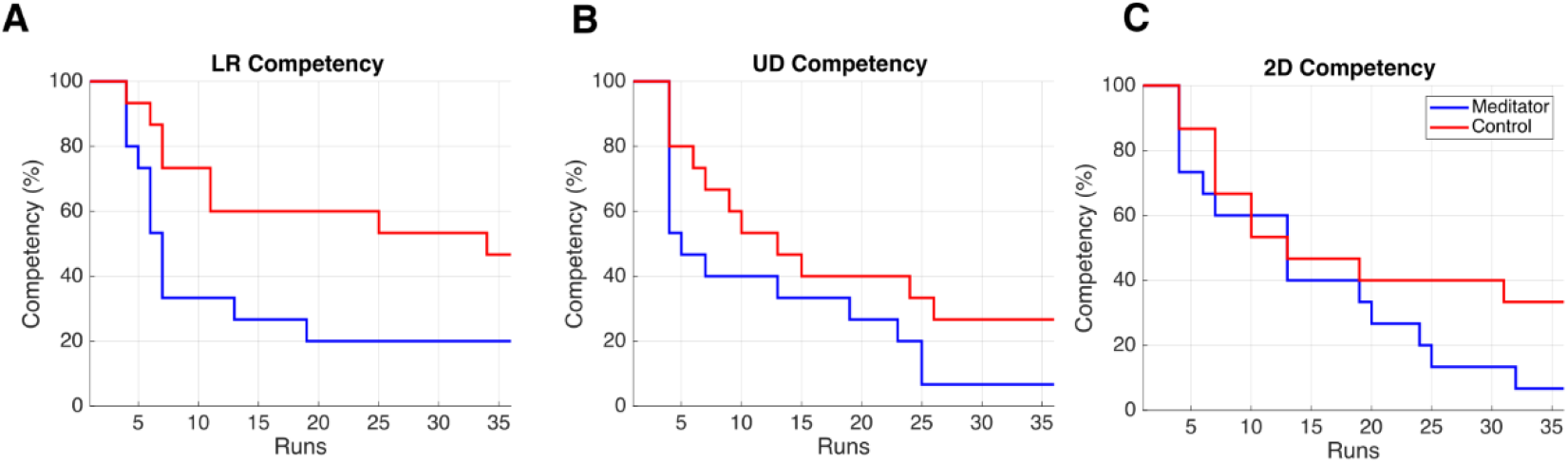
Competency curves for **(A)** LR, **(B)** UD and **(C)** 2D tasks. Each session has 6 runs for one task, accounting for 36 runs in total throughout the 6 training sessions. The numbers are percentage of subjects not meeting competency thresholds.

There are two observations from this plot. First, after six sessions of learning, the percentage of BCI inefficient subjects appears to be lower in meditators. The percentage of BCI inefficient subjects are 20% (46%), 6% (26%) and 6% (33%) for meditators(controls), in LR, UD and 2D tasks, respectively. Therefore, in all three tasks, meditators indeed had numerically less BCI inefficient subjects after 6 sessions or 36 runs of learning, but Chi-squared tests did not reveal significance for the proportion of BCI inefficiency between two groups (X^2^(1, N = 30) = 2.4, 2.16, 3.33, p = 0.12, 0.14 and 0.06 for LR, UD and 2D, p < 0.05). Second, the speed of learning, the LR and the UD plot showed a steeper decline during the initial 6 runs, i.e. the baseline session. This means that the learning speed of meditators appears to be faster than control subjects. Besides, while previous studies showed that BCI learning occurs in a session by session base (Meng et al., 2016), our results showed that learning could also occur within a 2-hour session. We also noticed that compared with 1D tasks (LR, UD), both groups in the 2D task showed a similar learning curve in the first 20 runs, i.e. in the first three sessions. After that, meditators showed a numerically better learning speed compared with control subjects. This observation is consistent with the previous group average performance in the sense that in UD within the 2D task, meditators had numerically larger improvement starting from the third session. In addition, it also shows that 2D control is indeed more difficult than 1D control, requiring more training time.

### 3.4 Group averaged topology during task

Figure 4 shows the LR and UD task fisher score topology (Perdikis et al., 2018) for meditators and controls. From the plot, a gradual increase of motor cortex area high alpha power could be seen in both groups, indicating that both groups were able to increase the contrast of two opposite conditions through voluntary motor imagery as learning progresses. However, this plot did not provide quantitative information regarding whether meditators had a higher baseline of C3 and C4 high alpha power, or if they exhibit better learning. To further investigate the effect of meditation experience on these quantities, we looked into the SMR predictor during the inter-trial resting state, and control signal during task execution.

**Figure 4.**
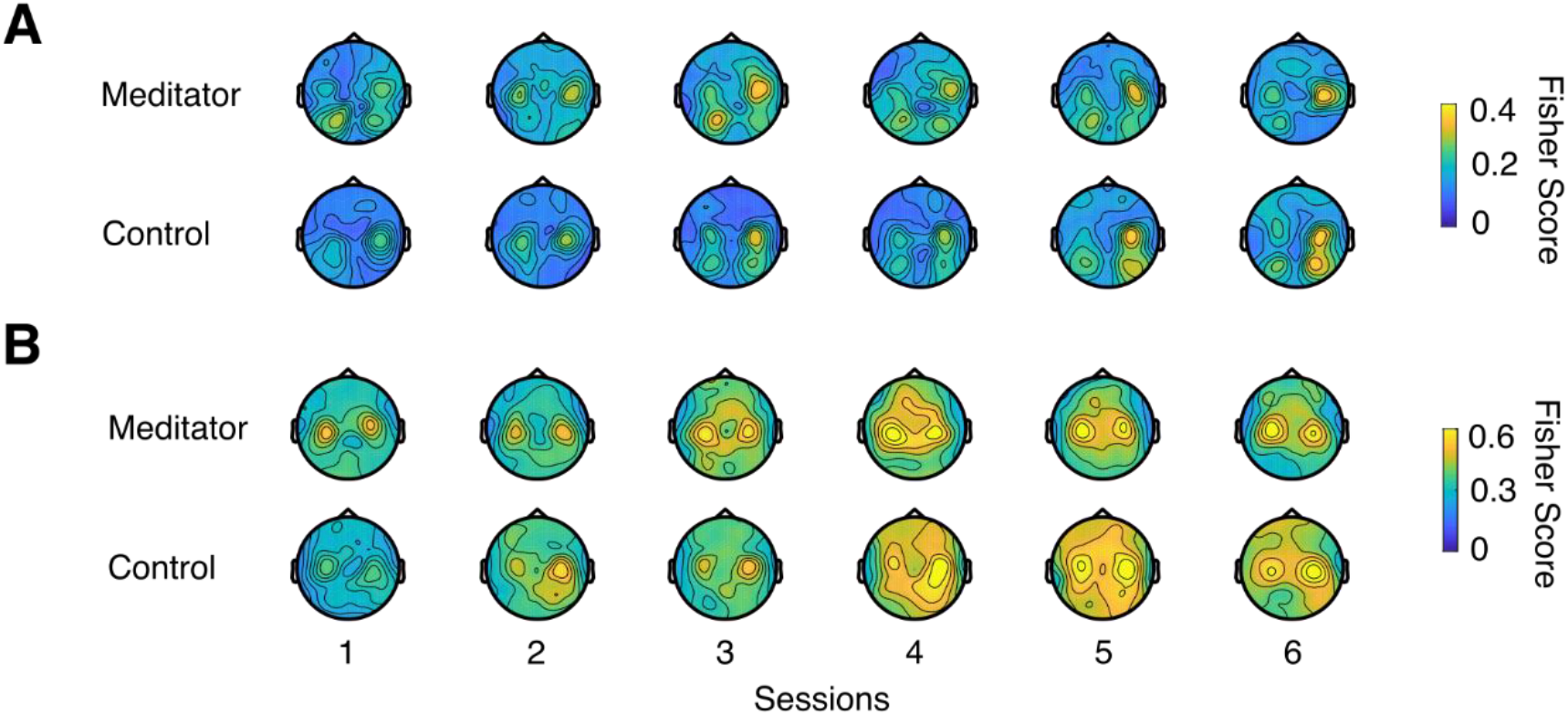
Fisher score topology for meditators and controls during the **(A)** LR and **(B)** UD task.

### 3.5 Neurophysiological predictor

Blankertz and colleagues (Blankertz et al., 2010) found that in the resting state power spectral density plot of C3 and C4 electrodes, the difference between mu rhythm peak and noise level baseline is a significant predictor of the BCI performance. Here we tried to investigate the difference in SMR predictor between meditators and controls. As shown in Figure 5 (A), we first fit a linear regression model between the SMR predictor and PVC. We found that in the LR task, the correlation coefficient between SMR predictor and PVC is r = 0.150 with a marginal significant p = 0.057, and in the UD task, r = 0.219 with p = 0.005. We noticed that there were outlier points for one subject that were far away from the population, therefore we also recalculated the correlation after removing this subject (Blankertz et al., 2010). After removing the outlier, in the LR task, the correlation coefficient was r = 0.385 with p < 0.001, and in the UD task, r = 0.268 with p < 0.001. Our correlation coefficient was smaller than that of Blankertz and colleague’s work (Blankertz et al., 2010). The difference might be due to the task design and subject variability. We next asked if experienced meditators have a better SMR predictor than controls. We found that the difference between the two groups was statistically significant (F(1,157) = 16.69, p < 0.001), but we did not observe the learning effect to be significant (F(1,157) = 2.2, p = 0.140, and there was no learning speed difference between groups (F(1,157) = 0.03, p = 0.859), as shown in Figure 5 (B).

**Figure 5.**
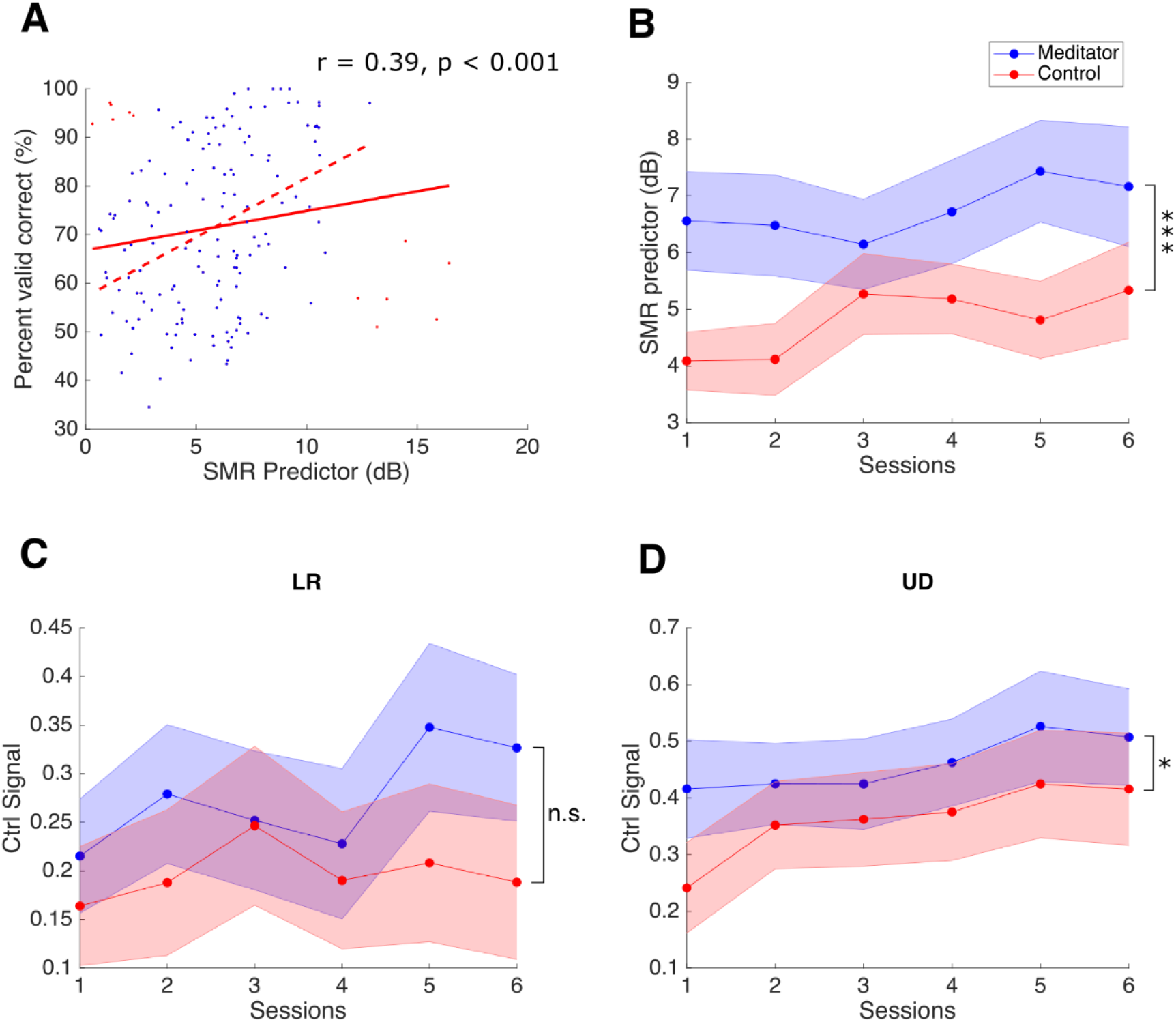
**(A)** Regression between SMR predictor and PVC, for LR task. The red line is the regression line, and red dashed line is the regression line after removing outlier (red points). The plot for UD task is similar and not shown. **(B)** Group averaged SMR predictor between meditator and controls, significance in the group effect is found. **(C, D)** Control signal learning as sessions go on for **(C)** LR and **(D)** UD.

### 3.6 Control signal baseline and learning

Given the behavior difference described in the previous section, the next question to ask is whether meditators exhibit better overall and learning of ΔCtrl Signal, defined as the control signal difference between two opposite motor imagery tasks (left versus right, up versus down). Figure 5(c)(d) shows the group averaged ΔCtrl Signal as sessions go on. For LR, before session 4, the two groups exhibited similar values, and starting from session 4, meditators had higher ΔCtrl Signal than controls, but the variance was high, and ANCOVA analysis did not reveal group, session, or the learning effect differences (F(1,176) = 3.34, 1, 0.5, p = 0.07, 0.32 and 0.48 for group, session or the learning difference). On the other hand, for the UD task, a numerical increase trend could be seen in both meditators and controls. The difference between the two groups was significant (F(1,176) = 4.19, p = 0.04), while the learning effect and learning rate difference was not (F(1,176) = 0.055 and 0.76).

## 4 Discussion

Reducing the training time and BCI inefficiency is critical in the application of SMR based BCI. While prior studies have tried to solve this problem from the ‘brain’ side of BCI by investigating the effect of meditation experience on SMR BCI learning, the relationship between these two is still not comprehensive. First, due to the large variability in the type and duration of meditation, more studies are needed to confirm the existence of such an effect. Second, it is still unclear whether and to what extend do meditators are better able to do more complex tasks than 1D control. Third, a more thorough investigation of the neurophysiological difference between these two groups is needed.

Our results provide insights into the effect of long-term meditation experiences on SMR based BCI. Concretely, we found that level of mindfulness is correlated with the SMR BCI performance in the UD task, and within the population who still have a margin to learn, experienced meditators had higher BCI performance compared with meditation naïve subjects. We also found that there were numerically fewer BCI inefficient subjects remaining after six sessions of learning. As for task complexity, we extended the control paradigm to a more complexed 2D cursor control task. We found a similar trend when separating the LR and UD tasks within the 2D control, that meditators had higher LR within the 2D performance than controls, we also found that although meditators and controls started at approximately the same level of UD within 2D performance, numerically, meditators had better learning and resulted in higher improvement than controls; Finally, neurophysiology analysis revealed that there is a significant difference between the SMR predictor and the UD control signal between two groups. In general, the statistical differences between these two groups mainly lie in the performance, we did not find the speed of learning to be significantly different between these two groups, which could be due to the reason that given meditators already have relatively higher performance, the room for improvement becomes smaller.

It should be noted that our experimental task is consistent with prior work done in the same lab (Cassady et al., 2014) in terms of the platform (BCI 2000) and 1D BCI task design. However, to the best of author’s knowledge, this study of comparing SMR based BCI performance between experienced meditators and controls has not been replicated. Our work was done in a different time, location, and subpopulation of experienced meditators and controls, yet we reveal results to some degree consistent with the previous work, supporting that differences exist between experienced meditators and controls in terms of SMR BCI control.

While the prior longitudinal study found an 8-week MBSR class mainly has effects on the UD trials (Stieger et al., 2020), our work showed that people with long term meditation experiences outperformed control subjects in both LR and UD tasks. This long-term meditation effect could be due to the plasticity introduced by meditation experience. For example, one of the main benefits of mindfulness meditation is enhanced attentional control (MacLean et al., 2010). In the SMR BCI, subjects are instructed focus on, or pay attention to the motor intention, which could regard as a specific type of attention control. Therefore, the prolonged meditation practices might serve as additional ‘training time’ and cause the meditator group to have enhanced BCI performance. Future work along this line should investigate if ordinary people are also able to improve SMR BCI control, apart from UD tasks (Stieger et al., 2020), with more extended meditation training.

An alternative explanation would be the pre-existing difference in the brain structure, personality, etc., for people who choose to meditate for years (Tang et al., 2015). In other words, the subpopulation who choose to meditate for years may have attributes that contribute to a successful SMR BCI control. Nevertheless, the research focusing on SMR BCI control ability for people with different characteristics is still limited, and future work on investigating the impact of these multidimensional and interrelated personal attributes might reveal more details of SMR BCI control.

Another concern regarding studying these two distinct groups is the effect of age. While we tried our best to find age-matched controls for the meditators, the meditators are on average 38.5 years old and controls are on average 24.8 years old years with a 13.7 years old difference. One might argue that the fact meditators being more senior might affect our conclusion. However, we did not find the correlation between age and performance to be statistically significant. Another evidence from prior study is that cortical physiology might decrease as people age (Roland et al., 2011). Therefore, the fact that meditator being more senior does not make our conclusions invalid.

## 5 Conclusion

In this study, we have examined the behavior and neurophysiological differences between experienced meditators and control subjects. We found that among subjects who still have a margin to learn, meditators outperformed control subjects in terms of averaged performance, SMR predictor and control signal (in the UD task). This finding has implications on enhancing the ‘brain’ side of SMR BCI and may help overcome the limitations of SMR BCI technology, such as long training time and BCI inefficiency.

## 7 Author Contributions

X.J. was involved in experiment conduct, data analysis and manuscript writeup. E.L. was involved in experiment conduct and manuscript review. J.S. was involved in study design, experiment conduct and manuscript review. C.G was involved in study design, subject recruitment and manuscript review. B.H. was involved in conception, study design, supervision, and manuscript review.

## 9 Acknowledgments

We would like to thank Dr. David Creswell for useful discussions, Chang Liu and Kristie Lindblom for assistance in subject recruitment, Olivia Fernau and Chalisa Udompanyawit for assistance in experiment conduct, and Dr. Haiteng Jiang for useful discussions.

## 11 Supplementary Material

**Table S1.** Group averaged performance from baseline and final session

| Identity \ PVC (%) |  | LR | UD | 2D |
| --- | --- | --- | --- | --- |
| Meditator | Baseline | 70.7 | 74.4 | 42.1 |
|  | Final | 79.6 | 79.2 | 50.4 |
| Control | Baseline | 62.8 | 62.9 | 36.5 |
|  | Final | 68.6 | 70.9 | 40.2 |

**Figure S1.**
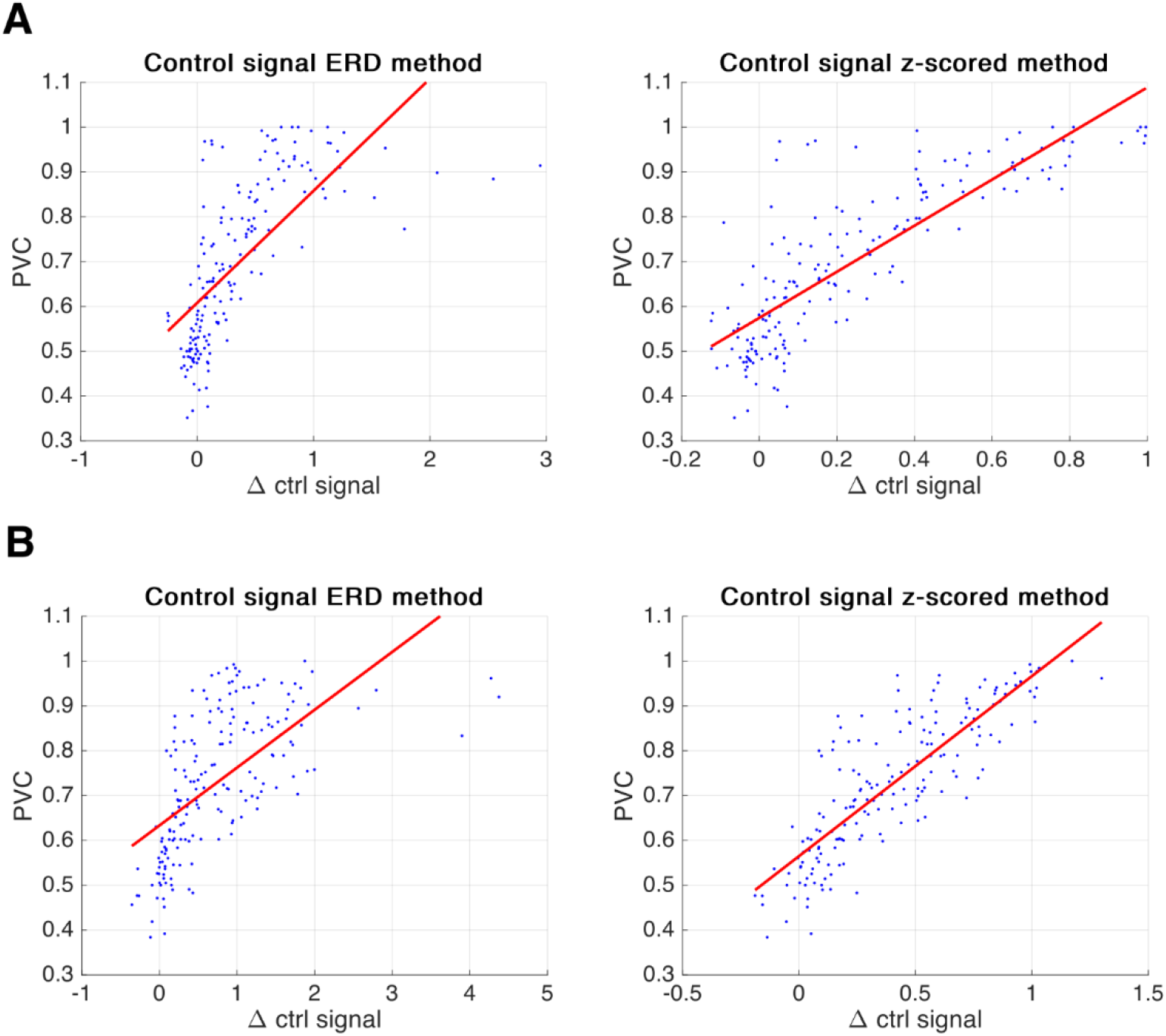
Comparison of two methods to compute the control signal during task execution. The traditional method to quantify how EEG band power changes during task execution is event-related desynchronization. Concretely, the control signal under the ERD definition would be band power normalized by the resting state alpha activity. Here we argue that the control signal using the z-score method would be a better metric by showing that it explains more performance variability. **(A)** in LR, the correlation coefficient for regression between ΔCtrl Signal and PVC was 0.69 and 0.83 in the ERD method and z-scored method, **(B)** for UD the it was 0.62 and 0.84, p < 0.05.

## 12 Data Availability Statement

The data presented here are available upon reasonable request from the corresponding author.

